# Exploring The Ability Of Machine Learning-Based Virtual Screening Models To Identify The Functional Groups Responsible For Binding

**DOI:** 10.1101/2023.04.29.538820

**Authors:** Thomas E. Hadfield, Jack Scantlebury, Charlotte M. Deane

**Affiliations:** Oxford Protein Informatics Group, Department of Statistics, University of Oxford, Oxford OX1 3LB, UK

## Abstract

Many recently proposed structure-based virtual screening models appear to be able to accurately distinguish high affinity binders from non-binders. However, several recent studies have shown that they often do so by exploiting ligand-specific biases in the dataset, rather than identifying favourable intermolecular interactions in the input protein-ligand complex. In this work we propose a novel approach for assessing the extent to which machine learningbased virtual screening models are able to identify the functional groups responsible for binding. To sidestep the difficulty in establishing the ground truth importance of each atom of a large scale set of protein-ligand complexes, we propose a protocol for generating synthetic data where the label of an example is assigned by a 3-dimensional deterministic binding rule. This allows us to precisely quantify the ground truth importance of each atom and compare it to the model generated attributions.

Using our generated datasets, we demonstrate that a recently proposed deep learning-based virtual screening model, PointVS, identified the most important functional groups with 39% more efficiency than a fingerprint-based random forest, suggesting that it would generalise more effectively to new examples.

In addition, we found that ligand-specific biases, such as those present in widely used virtual screening datasets, substantially impaired the ability of all ML models to identify the most important functional groups.

We have made our synthetic data generation framework available to facilitate the benchmarking of new virtual screening models. Code is available at https://github.com/tomhadfield95/synthVS.

## Introduction

The drug discovery process is difficult and time consuming, with a recent study finding that the median time to develop a new drug was 8.3 years, at an average cost of $985 million.^1^ There is therefore a need to develop novel techniques which can help design medicines more quickly and cheaply.

Fueled by their successes in a broad range of domains (e.g.^2–4^), there has been substantial recent interest in the development of machine learning (ML) algorithms to help accelerate the drug discovery process; recent examples include highly accurate protein structure prediction algorithms,^4^ generative models which propose target-specific libraries for compound design^5^ and tools for automated synthesis planning for synthetically challenging molecules.^6^

The long-term hope for these algorithms is that it will one day be possible to build an end-to-end *in-silico* drug-discovery platform which, following the identification of a target, could produce a series of high affinity binders without human involvement. A key component of such a platform would be the ability to computationally predict whether a given ligand is likely to bind to the target with high affinity, as this would allow the prioritisation of promising compounds from a large library of molecules.

Traditional virtual screening models (e.g.^7,8^) estimate the binding affinity of a protein-ligand complex as a weighted sum of physics-based terms (e.g. van der Waals contributions, hydrogen bond scores etc.), with solved protein-ligand complexes used to estimate the model weights. These methods are dependent on a set of hand-crafted features being fed into a model; subsequent approaches have sidestepped this requirement by converting an input molecule to a fingerprint representation and using machine learning algorithms to automatically approximate non-linear relationships between features, such as random forests^9^ or neural networks.^10^

Inspired by the remarkable ability of deep learning models to capture important spatial information in other fields, the most recent set of virtual screening models (e.g.^11–13^) have tended to use deep-learning architectures, representing the protein-ligand complex either as a graph or as a 3D image. It is hypothesized that such models are better able to identify important protein-ligand interactions and will be more generalisable to novel targets than fingerprint-based models or models which depend on a set of hand-picked features.

Despite their ability to capture important spatial information, recent studies^12,13^ have shown that fingerprint-based models (e.g.^10^) illustrated comparable or better predictive performance than deep learning models, calling into question whether deep learning models are actually able to use their learned representations to identify important intermolecular interactions. Moreover, a study by Chen et al^14^ raised concerns surrounding the susceptibility of deep learning-based models to ligand-specific biases, demonstrating that removing all protein information from a deep learning-based virtual screening model did not degrade its performance. Their findings indicated that the model was not capturing important spatial information but rather learning to classify examples based on ligand-specific biases. In a recent study by Volkov et al. ^15^, the authors trained a series of message passing neural networks with a variety of different inputs. They found that whilst the inclusion of protein-specific information aided the model’s predictive performance compared to a model trained solely on ligand-specific features, the explicit featurisation of protein-ligand interactions did not improve performance compared to the model trained on ligand-specific and protein-specific features. The authors argued that this suggested that the model was unable to learn to identify the underlying biophysical interactions responsible for binding and instead learned to classify examples based on distributional differences in the training data.

An approach to incentivise models to learn to use intermolecular interactions was described by Scantlebury et al.^16^ They proposed a data augmentation strategy where additional decoy examples were derived by taking an active protein-ligand complex from the training set and randomly rotating and translating the ligand. The augmentation procedure forced the convolutional neural network to use the protein-specific information when making decisions, illustrated by the degraded performance of the model when protein-specific information was removed.

To investigate the extent to which ML algorithms are able to accurately assign importance to individual atoms when making a prediction, several recent works^17–19^ have proposed synthetic datasets which labelled ligands as ‘active’ if they contained a pre-defined molecular substructure. Using an attribution technique, such as Integrated Gradients, ^20^ it is then possible to compare the model-assigned atom importances to the ground truth atom labels to assess whether the atoms which comprised the predefined substructure were identified as the most important atoms. While these studies provided valuable insights into the ability of ML algorithms to identify important functional groups, they did not assess whether the ML algorithms were able to capture important spatial information or identify intermolecular interactions.

Whilst several authors have used attribution techniques on real-world data to uncover important functional groups, ^16,21,22^ it is often difficult to ascertain the precise contribution of each atom in an experimentally obtained proteinligand complex. Combined with the difficulty in manually curating a large-scale test set, it is currently infeasible to objectively assess the attribution performance of ML algorithms on real-world virtual screening tasks.

To address this, we propose a protocol for generating a synthetic dataset which mimics the mechanics of protein-ligand binding. As the label of each example is determined by a known deterministic binding rule, we can precisely specify which functional groups, if any, in the ligand are responsible for binding. Although our deterministic binding rule is considerably simpler than the mechanics of real-world protein-ligand recognition, it allows us to assess whether an ML algorithm is able to capture important spatial information and use it when making predictions.

Using our synthetic dataset, we first quantified the ability of a fingerprint-based virtual screening model to correctly identify important functional groups, and investigated the effect of changing the model’s parameters on its attribution performance. We then investigated the effect of ligand-specific biases on the fingerprint-based models, both in terms of predictive accuracy and attribution performance. Finally we compared the performance of the fingerprint-based models to a recently proposed Equivariant graph neural network, PointVS.^22^

We found that although all models were able to accurately predict binding in the presence of ligand-specific biases, their ability to attribute binding to the correct functional groups was substantially degraded, indicating they were less able to generalise than models which were not susceptible to ligand-specific biases. We also found that the attribution performance of the fingerprint-based models was heavily dependent on the parameters used to define the fingerprint, and that they were less able to identify the most important functional groups compared to the EGNN method, PointVS. These findings illustrate the importance of investigating the reasons behind high predictive performance and the utility of our synthetic approach as a benchmark for model generalisability.

## Methods

It is not easily possible to precisely experimentally quantify the extent to which an intermolecular interaction contributes to proteinligand binding on a large scale on real-world data. Therefore we propose two protocols for generating synthetic data where the contribution of any atom can be computed exactly. Whilst the synthetic data we generate gives only a coarse-grained approximation of realworld protein-ligand binding, it nevertheless allows us to assess the ability of virtual screening models to utilise important spatial information. We are also able to ensure that our synthetic datasets are free of ligand-specific bias and other extraneous factors which may inflate a model’s predictive performance on a test set while degrading its generalizability to novel targets. This allows us to quantify the effect of such real-world factors on model generalizability.

### Generating a synthetic protein-ligand complex

We defined a “synthetic protein” to be the set {(*x*_*i*_, *y*_*i*_, *z*_*i*_, *t*_*i*_)|*i* = 1, …, *m*}, where each element, which we call a “synthetic residue”, comprises 3D coordinates (*x*_*i*_, *y*_*i*_, *z*_*i*_) and an associated type, *t*_*i*_. After specifying a ligand with a 3D conformation, we constructed a synthetic protein as follows:

- We first defined a box around the ligand. To obtain the *x*-axis of the box we identified the minimum and maximum *x*-coordinates, *x*_*min*_ and *x*_*max*_, over all ligand atoms and defined the *x*-axis as [*x*_*min*_ *−* 5 Å, *x*_*max*_ + 5 Å]. The *y*-axis and *z*-axis were obtained in the same way.
- We then sampled a number of coordinates uniformly within the box, such that the density of points was invariant of the box size. That is, we sampled *m* points, where *m* = *a*_*coef*_ × *b* for box area *b*.
- For each set of coordinates, we randomly sampled an associated type to create a synthetic residue.
- If any synthetic residue was within 2 Å of a ligand atom, it was deleted.
- The synthetic residues were filtered further so that no two synthetic residues are within 3 Å of each other.
- To reduce the risk of inducing ligand-specific bias dependent on the number of functional groups present in a ligand, we sampled the number of synthetic residues, *n*_*res*_, as *n*_*res*_ = *n*_*ops*_*/n*_*lig*_, where *n*_*ops*_ is a constant and *n*_*lig*_ was the number of ligand functional groups which can interact with a protein (which varies according to the generative process used to generate the synthetic protein, see below).

### Polar generative process

The type of each synthetic residue determines which ligand functional groups it is able to interact with.

In the Polar dataset we restrict the synthetic residue types to “Hydrogen Bond Acceptor” (HBA) and “Hydrogen Bond Donor” (HBD). We used RDKit^23^ to determine whether each ligand atom was a Hydrogen bond donor or acceptor and we define a ligand donor or acceptor to interact with a synthetic residue if:

- Their types match; e.g. the synthetic residue has type HBA and the ligand atom is a Hydrogen Bond Acceptor, and
- The distance between the synthetic residue and ligand atom is below a specified threshold. For all experiments this was set at 4 Å.

For a given synthetic protein-ligand complex, if any ligand atom interacts with a synthetic residue, we say the complex is active, otherwise it is inactive. An example of a synthetic protein-ligand complex generated using the Polar generative process is shown in Figure 1.

**Figure 1:**
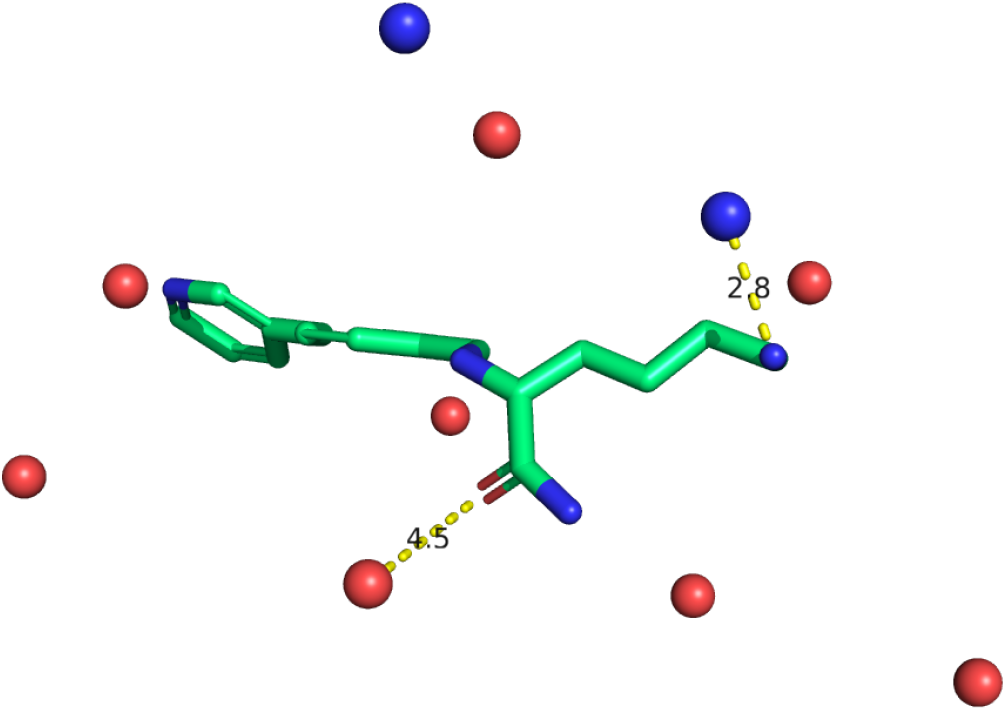
Example of synthetic protein-ligand complex. Green ligand atoms represent Carbons, red atoms denote Oxygens and blue atoms denote Nitrogens. The blue spheres represent synthetic residues with type ‘Hydrogen Bond Donor’ (HBD), whilst the red spheres represent synthetic residues with type ‘Hydrogen Bond Acceptor’ (HBA). The amine group is 2.8 Å away from a synthetic residue with type HBD; as the distance is less than the 4 Å specified by the deterministic binding rule, we consider this example to be active. While the carbonyl is 4.5 Å away from the nearest synthetic residue of with type HBA (and therefore does not interact with it), under the Polar generative process only a single interaction is needed for binding.

### Contribution-based generative process

While the synthetic protein-ligand complexes generated by the Polar generative process only require a single interaction to be classified as active, in practice a ligand typically needs to make several different interactions in order to bind with high affinity. We therefore propose a second binding rule which assigns a score to each synthetic residue-ligand atom pair and classifies the complex depending on the cumulative score. In the Contribution dataset we extend the synthetic protein to include “Hydrophobic” synthetic residues as well as the HBA and HBD synthetic residues present in the Polar dataset. The synthetic protein is generated in the same way as when using the Polar generative process, with each synthetic residue type sampled at uniform from the set *{HBA, HBD, Hydrophobic}*.

To label a synthetic protein-ligand complex, we consider synthetic residue-ligand functional group pairs where the ligand functional group has type “Acceptor”, “Donor” or “Hydrophobe”, assigned by RDKit.^23^ If the synthetic residue and ligand functional group do not have matching types, we define their interaction score as 0, otherwise their interaction score is calculated as a non-linear function of the interaction type and the euclidean distance between the synthetic residue and ligand atom (Figure 2). The sum of all pairwise interaction scores is calculated, and we label the example as “active” if the summed interaction score exceeds a pre-specified threshold. For all experiments in this work, we used a score threshold of 4.

**Figure 2:**
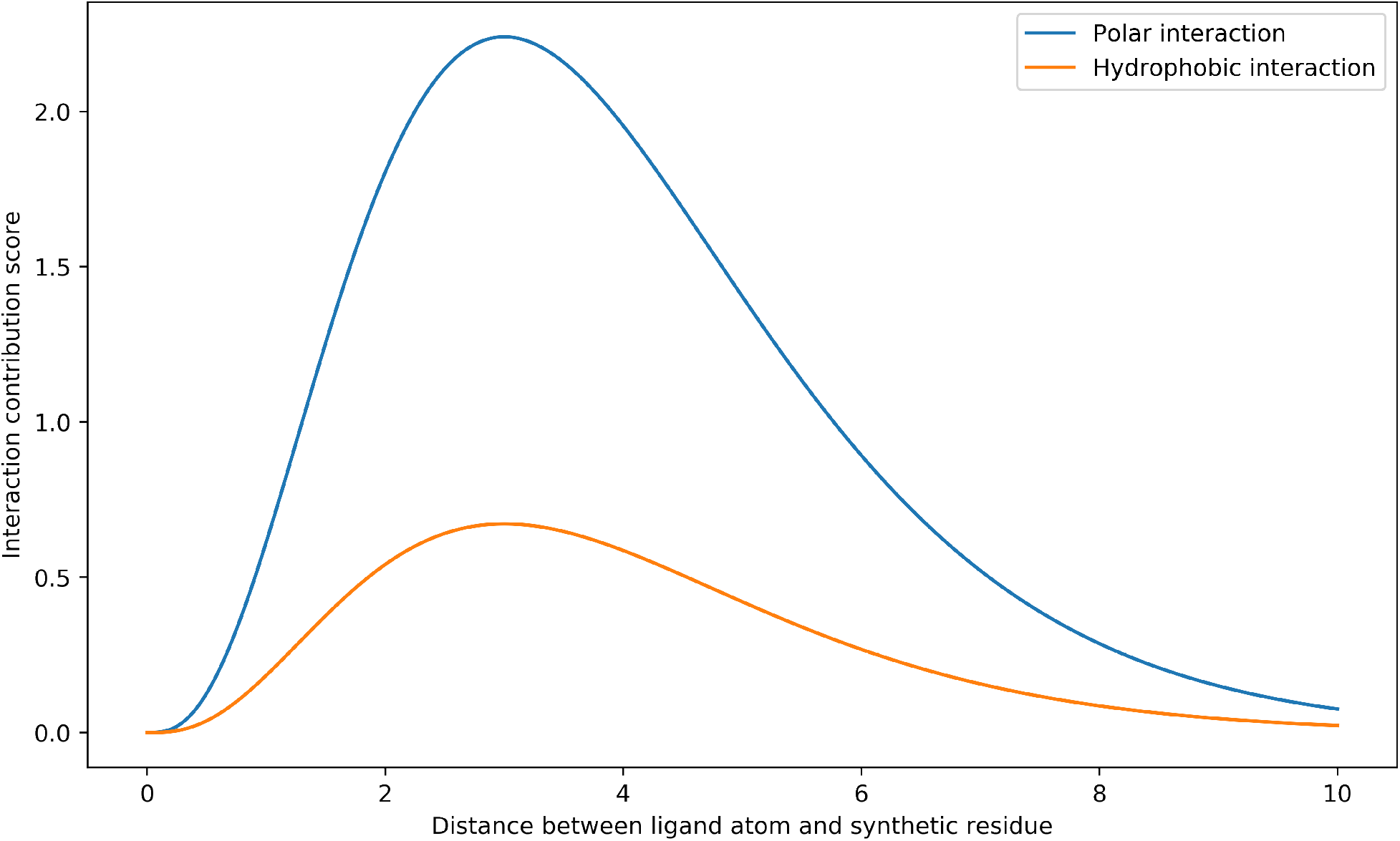
Non-linear functions used to determine the interaction score of a synthetic residueligand atom interaction; both functions are proportional to the Probability Density Function of a Gamma(4, 1) distribution. Separate functions are used for Polar interactions and Hydrophobic interactions, making it more difficult for the model to learn the deterministic binding rule.

### Machine learning algorithms

We used two different ML algorithms to classify examples as active or inactive. The first model was a random forest (RF), which took either Morgan fingerprints^24^ or Protein-Ligand Extended Connectivity fingerprints (PLECs)^25^ as input. We were able to represent our synthetic protein-ligand complexes as PLECs without needing to modify the Open Drug Discovery Toolkit^26^ (oddt) codebase.

#### Morgan Fingerprints

Morgan fingerprints are a vector representation of a molecule, calculated in an iterative fashion by first assigning an identifier to each atom and then updating the identifier to incorporate information from the identifiers of the atom’s neighbours. This updating process is repeated a number of times so that information about atoms which are not immediate neighbours of an atom can be included in its identifier; in this work, all Morgan fingerprints are calculated using the RDKit^23^ implementation with a radius of 2.

Morgan fingerprints, by design, do not incorporate any spatial information or any information about the target. In this work, the models trained using Morgan fingerprints serve as a baseline to assess whether a model can attain strong predictive accuracy using solely ligand-based features. Predictive performance that was substantially better than random would suggest that significant ligand-specific biases were present in the dataset, whereas close-to-random predictive performance would indicate that the active and inactive ligand sets were drawn from approximately the same population.

This allows us to quantify the extent to which a model might be susceptible to ligand-specific biases, as we would not expect models without access to information concerning the protein to be able to learn the deterministic binding rules and therefore better-than-random predictive accuracy would likely be due to biases in the training set.

#### PLEC Fingerprints

PLEC fingerprints encode important spatial information as follows: First, all ligand atom-protein atom pairs which are closer together than a specified cutoff are identified, and integer radii, *r*_*lig*_ and *r*_*prot*_, are specified for the ligand and protein respctively. (which need not be the same length). For each qualifying ligand atom-protein atom pair, substructures containing atoms which are up to *r*_*lig*_ or *r*_*prot*_ atoms away from the ligand or protein atom are identified and encoded in the fingerprint using a hashing algorithm. All PLEC fingerprints were computed using the oddt^26^ implementation with a ligand radius of 3 and a protein radius of 0 (as the synthetic residues are represented as a single, unconnected atom).

As ligand atom-protein atom pairs which are within a specified threshold are explicitly encoded within the fingerprint, we would expect models trained using PLECs to perform strongly on the Polar tasks when the PLEC distance cutoff closely matches the distance specified by the deterministic binding rule. However, as PLEC doesn’t encode any more detailed spatial information, we would expect that it would be unable to learn the non-linear deterministic binding rule used by the Contribution generative process.

#### Model naming

We refer to models trained using a Morgan fingerprint as `RF_Morgan`. Models trained using PLEC fingerprints are referred to `RF_PLEC`. `RF_PLEC_n`, where `n` is a positive real number, refers to an RF model trained with a specific PLEC distance cutoff.

### Equivariant Graph Neural Network (EGNN)

We compared the performance of the `RF_PLEC` models to a recently proposed deep learning-based approach,^22^ “PointVS”. PointVS is based on the E(n)-Equivariant Graph Neural Networks proposed by Satorras et al.^27^ In contrast to the fingerprint-based `RF_PLEC` models, PointVS takes as input the 3D coordinates of protein and ligand atoms, in addition to a one-hot encoding of the atom type. When constructing the input graph, where each node is an atom, two nodes are connected by an edge according to the following rules:

- two ligand atoms are connected by an edge if they are within 2 Å of each other.
- two protein atoms are connected by an edge if they are within 2 Å of each other.
- a protein atom and a ligand atom are connected by an edge if they are within 10 Å of each other.

For intra-molecular edges, the 2 Å cutoff connects atoms which are covalently bonded, whilst the inter-molecular edges connect atoms which might potentially interact. Whilst PointVS enforces a protein atom-ligand atom distance cutoff in the same way as PLEC, its purpose is to reduce the dimensionality of the input graph by not connecting atoms which are too far away from each other to interact with non-negligible strength. PointVS is unlikely to be able to utilise its distance cutoff to discover high energy interactions, as all protein and ligand atoms are connected if they are within 10 Å. It must instead learn to identify important interactions from the atomic coordinates and atom types. Scantlebury et al. ^22^ also applied a further distance cutoff, where any receptor atom which was not within 6 Å of any ligand atom was ignored; this reduced the dimensionality of the input graph by ignoring residues which were not part of the binding pocket. As we constrained each synthetic protein to be within a box defined as [*x*_*min*_*−*5, *x*_*max*_+5]×[*y*_*min*_*−*5, *y*_*max*_+5]× [*z*_*min*_*−*5, *z*_*max*_ + 5], where *x*_*min*_ was the smallest ligand atom x-coordinate and the other values were defined similarly, the vast majority of synthetic residues would be within 6 Å of at least one ligand atom and so this cutoff should have minimal impact on the performance of PointVS.

### “ZINC” dataset

Before exploring the effect of ligand-specific bias on model attribution performance, we constructed a baseline dataset to assess the ability of different ML algorithms to learn the deterministic binding rules outlined above. We obtained a set of 10k ligands from ZINC^28^ and used the Polar and Contribution generative processes to generate two synthetic datasets (“Polar_ZINC” and “Contribution_ZINC”).

We sampled 500 examples from each dataset to serve as a test set, and used the remaining examples to train each model. We trained 8 different `RF_PLEC` models, varying the PLEC distance cutoff by 0.5 Å from 2.5 Å to 6 Å. To train PointVS, we used the default hyperparameters outlined by Scantlebury et al. ^22^.

### Inducing ligand-specific bias

As mentioned above, Chen et al. ^14^ showed that the presence of ligand-specific biases allowed virtual screening models to disregard the provided protein-specific information and still achieve strong predictive accuracy. The models were able to do this by constructing a decision rule which classified examples based on 2D ligand features rather than intermolecular interactions. We hypothesized that a dependence on ligand-specific biases would severely degrade the ability of a virtual screening model to identify important functional groups.

To explore the effect of ligand-specific bias on attribution performance, we constructed a set of synthetic protein-ligand complexes where each ligand was taken from the Directory of Useful Decoys - Enhanced (DUD-E).^29^ Sieg et al^30^ showed that ML algorithms were able to exploit distributional differences between the actives and inactives in DUD-E to achieve inflated predictive accuracy. We selected the five DUD-E targets containing the most ligands and used the ligands to construct five synthetic datasets. For each ligand, we extracted its true label from DUD-E and generated a synthetic protein using the Polar generation process outlined above. If the synthetic protein-ligand complex had a different label than the true label assigned to the ligand, we generated a new synthetic protein and recalculated the label until the true label and the label of the synthetic protein-ligand complex matched. If, after generating 100 synthetic proteins we were unable to attain the true label, we discarded the ligand. We used the same approach to derive an additional five synthetic datasets from the five LIT-PCBA^31^ targets with the most ligands.

By constructing the synthetic virtual screening datasets in this way, an ML algorithm would be able to classify examples by using the spatial information in the synthetic protein-ligand complexes and/or by learning ligand-specific biases, mimicking the choice offered to an algorithm trained on the real DUD-E and LIT-PCBA datasets.

### “PDBBind” dataset

We used ligands from the PDBbind set^32^ (v. 2019) to construct two external test sets, one using the Polar generative process and one using the Contribution generative process. To reduce the risk of inflated performance due to overfitting, any ligand with a Tanimoto similarity of more than 0.8 to any ligand in any of the above training sets was discarded and we also discarded any very small molecules (fewer than 15 heavy atoms). Of the remaining ligands, we randomly selected 500 and generated a synthetic protein-ligand complex from each. As the purpose of the test set was to assess the ability of the models to identify functional groups which were responsible for binding, we ensured that each example was “active” by resampling the synthetic protein for each ligand until it was active according to the deterministic binding rule.

### Ranking the importance of ligand atoms

Following Hochuli et al, ^21^ we used atom masking to rank the importance assigned to a ligand atom by a particular model. For the *i*^*th*^ atom in a molecule, *m*, we calculate the masking score as *s*_*i*_ = score(*m*) *−*score(*m\i*), where score(*m*) is the prediction given by the model for *m* and *m\i* denotes the molecule *m* where the *i*^*th*^ atom has been deleted. For the `RF_PLEC` models, we delete an atom by replacing it with a dummy atom, and for PointVS we delete the corresponding node from the input graph. As higher scoring molecules are classified as active examples, masking assigns a high level of importance to atoms whose omission drastically reduces the model’s confidence that an example has an active label. Whilst Scantlebury et al. ^22^ used an attention mechanism to score the relative importance of different atomic interactions, we used masking for the experiments in this paper as it can be used to generate attributions for any predictive model, allowing a closer comparison between different models.

### Evaluation metrics

For datasets generated using the Polar process, where each atom either contributes to binding or is not involved at all, we would hope that an attribution method would give the highest rank to atoms involved in binding, allowing users to identify the most important atoms by their attribution scores. We propose an ‘Attribution AUC’, where a ranking of atoms which places all binding ligand atoms at the top receives a score of 1, a ranking which places all binding ligand atoms at the bottom receives a score of 0, and all other rankings receive a score according to the following heuristic:

We define a ‘change’ to be the transposition of two adjacent rows in a dataframe sorted by the model attributions. We calculate the number of changes required to rank the ligand atoms correctly (all binding atoms ranked at the top), and the number of changes required to rank the ligand atoms correctly in the worst case scenario (all binding atoms ranked at the bottom).

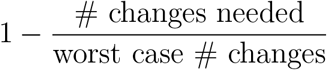

For datasets generated using the Contribution process, we calculated the Spearman’s Rank Correlation Coefficient between the true ligand atom contributions and the model attributions for each example. To assess the extent to which the models were able to identify the most important atoms, we introduce two metrics:

- Mean Above-Threshold Ranking (MATR): For a specified score threshold, *t*, we compute the average rank (assigned by the model attributions) of all atoms whose true contribution was greater than *t*. A low MATR implies that a model can successfully identify important atoms; we compute the MATR for score thresholds between 0 and 3.
- Relative Efficiency of Ranking (RER): We compare the atomic rankings derived from the model attributions to the ”perfect” ranking derived from the ground truth atomic contributions. We define:

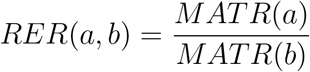

where *MATR*(*a*) is the MATR score attained by the model *a*. For the RER score, we calculate the MATR scores by calculating the average model-assigned rank of the top decile of atoms.

In addition to the above metrics which assess the ability of a model to correctly identify which atoms were responsible for binding, we also computed the accuracy and area under the Precision-Recall curve (AU PRC) attained by the models on the held out test sets.

## Results and Discussion

In order to quantify the extent to which different ML algorithms were able to accurately identify important intermolecular interactions, we tested the algorithms on synthetic datasets using deterministic binding rules (see Methods).

### Ligand-only model performance

In line with previous work,^14,16^ we trained several models where no synthetic protein-specific information was provided to the model (`RF_Morgan`, see Methods). The performance of these models allowed us to assess whether a dataset exhibited any ligand-specific biases, as the deterministic binding rule was dependent on both ligand and synthetic-protein.

All `RF_Morgan` models trained on DUD-E datasets attained substantially better-than-random predictive accuracy (Table S1). This was also true for all LIT-PCBA ligands if balanced numbers of actives and decoys were used (Table S1). This suggests that `RF_Morgan` models trained on DUD-E or LIT-PCBA datasets were able to exploit ligand-specific information to make accurate predictions. By contrast, when trained on the Polar_ZINC dataset, the `RF_Morgan` model attained a predictive accuracy of 0.52, indicating that without target-specific information a Random Forest using only ligand information was unable to accurately classify the examples (Table 1).

**Table 1:**
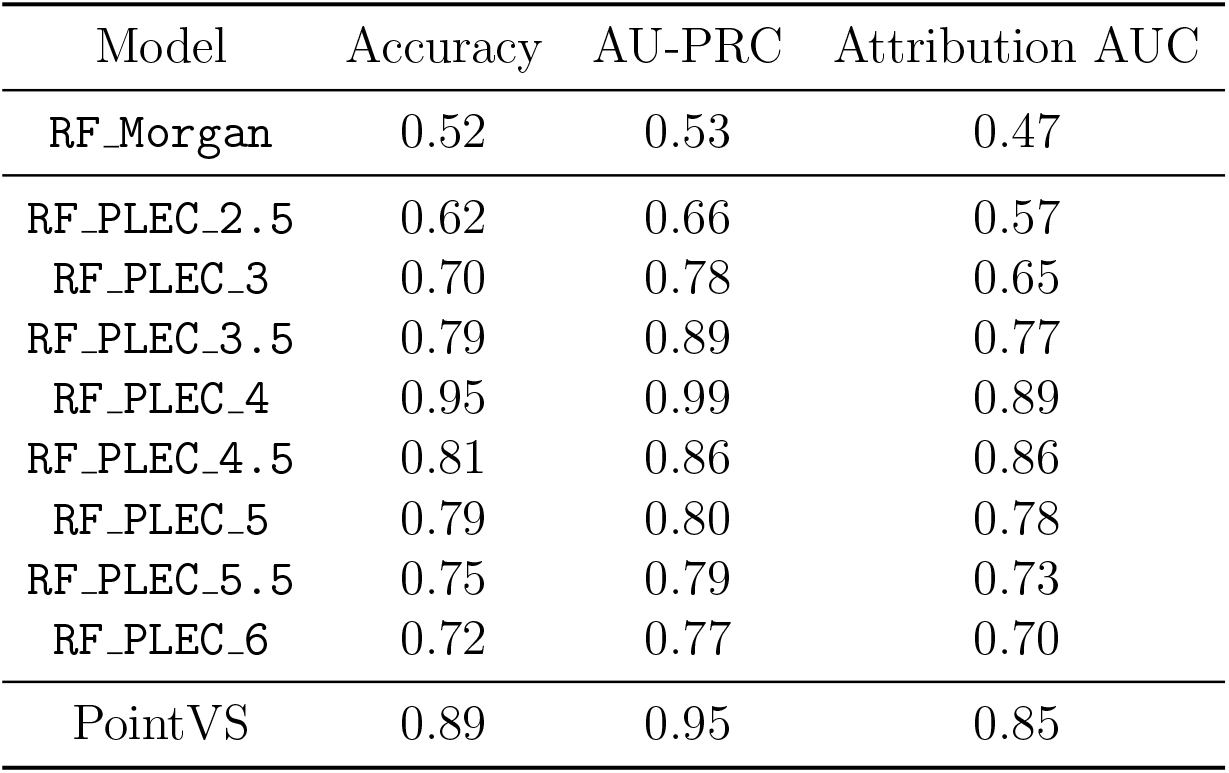
Performance of different `RF_PLEC` models when trained on the Polar_ZINC dataset. Accuracy denotes the proportion of correctly classified examples on the Polar_ZINC test set, AU-PRC denotes the area under the Precision-Recall curve on the Polar_ZINC test set, and Attribution AUC reflects the ability of a model to correctly identify the ligand atoms responsible for binding on the PDBBind test set (see Methods). The best performing model was `RF_PLEC_4`, which uses the same distance cutoff as the Polar deterministic binding rule.

### Performance of RF_PLEC models (ligand + protein fingerprints) on Polar_ZINC dataset

Having validated that our Polar_ZINC dataset did not contain trivial ligand-specific biases, we built `RF_PLEC` models that include the synthetic proteins in the training process (see Methods). To assess how sensitive the `RF_PLEC` models were to the choice of PLEC distance cutoff, we trained eight distinct `RF_PLEC` models, incrementing the PLEC distance cutoff by 0.5 Å between 2.5 Å and 6 Å. We found that all `RF_PLEC` models attained a substantially better predictive accuracy than the `RF_Morgan` model on the Polar_ZINC dataset (Table 1), indicating that the inclusion of protein information into the model enabled the random forest to better approximate the deterministic binding rule. Unsurprisingly, the `RF_PLEC_4` (the same cutoff as the deterministic binding rule) model obtained the highest predictive accuracy (Table 1).

The `RF_PLEC` models exhibited a similar trend in terms of Attribution AUC (which assesses the ability of a model to assign high ranks to active ligand atoms), with the `RF_PLEC_4` model most often correctly identifying the atoms responsible for binding (Table 1). We observed that performance degraded substantially as the PLEC distance cutoff diverged from 4 Å, suggesting that the `RF_PLEC` models were highly dependent on the precise specification of the PLEC distance cutoff.

### Sensitivity of RF_PLEC models to distance cutoff

To better understand the relationship between the PLEC distance cutoff and attribution performance, we consider a synthetic protein-ligand complex in detail. The original complex (shown in Figure 3a) is labelled as active and has a single synthetic residue-ligand atom pair with complementary type and a distance below the deterministic binding threshold. We perturbed the synthetic residue to take a range of locations between 1.5 Å and 10 Å away from the matching ligand atom (Figure 3b). We featurized each protein-ligand complex using PLEC and used masking to determine the importance of each ligand atom and calculated the rank of the ligand atom involved in binding. Figure 3c shows the relationship between synthetic residue-ligand atom distance and the rank assigned to the binding ligand atom by the highest performing `RF_PLEC` models (RF_PLEC_4, RF_PLEC_4.5 & RF_PLEC_5). Figure 3c shows that the `RF_PLEC` models assign a high level of importance to the key ligand atom when the synthetic residue-ligand atom distance is less than the respective PLEC distance cutoff, and assigns a reduced level of importance when the distance is greater than the PLEC distance cutoff. In particular, when using a PLEC cutoff threshold of 4.5 or 5 Å, the ligand atom is assigned a high rank when the synthetic residue-ligand atom distance is greater than the true binding threshold but less than the PLEC distance cutoff. This demonstrates the sensitivity of the `RF_PLEC` models to the exact specification of its distance cutoff; it is not able to encode a precise level of spatial information. This suggests that models trained using PLEC or based on distance-based cutoffs are unable to retain fine-grained spatial information, which may limit their usefulness for work with real-world protein-ligand complexes where different interactions can take place at different distances.

**Figure 3:**
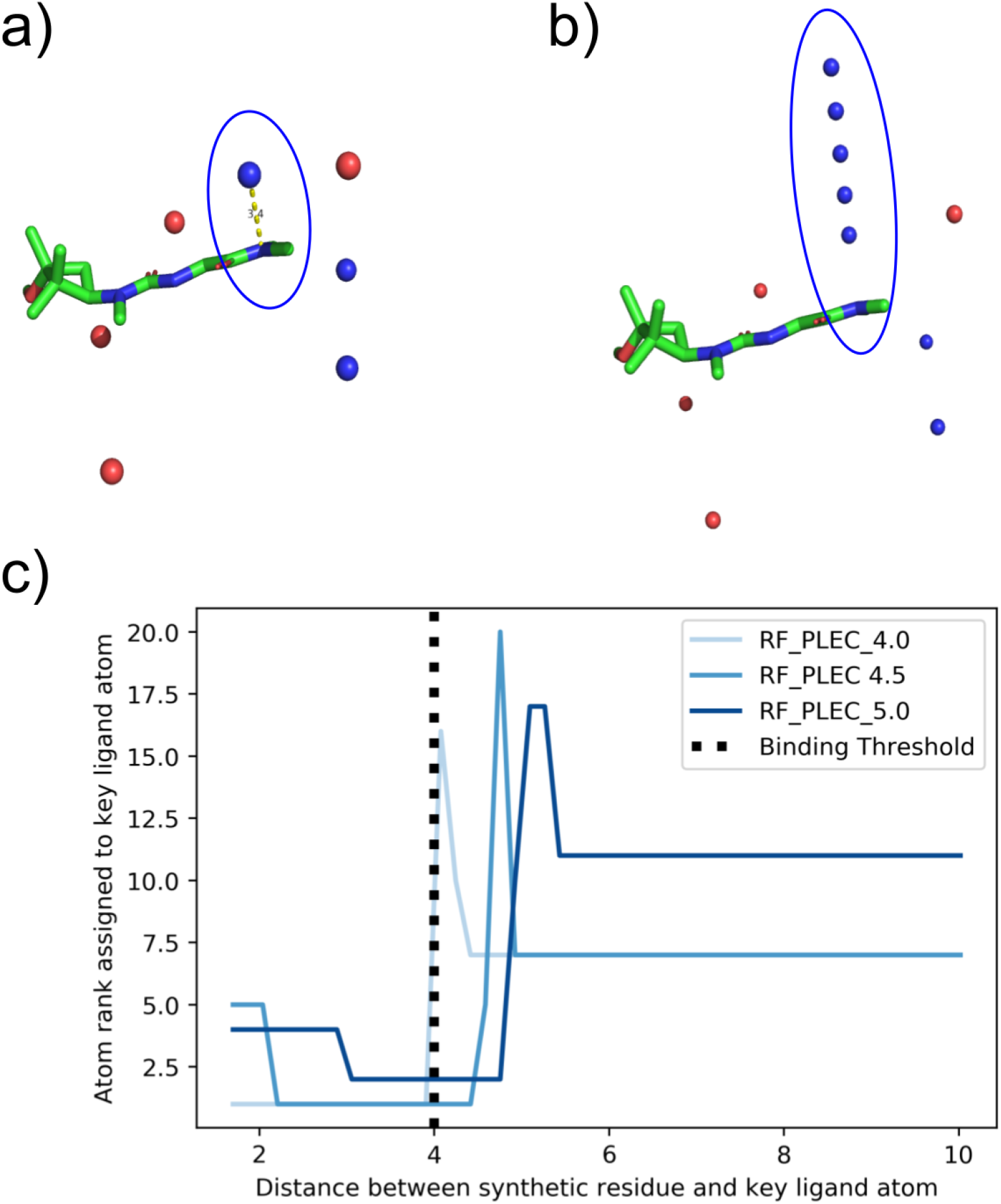
Case study illustrating the effect of changing the PLEC distance cutoff on model attributions. a) The original synthetic protein-ligand complex generating using the Polar generative process, labelled as active as a result of the circled interaction. b) Perturbed examples. We generated 50 synthetic protein-ligand complexes, which were all identical apart from the HBD synthetic residue contained in the circle, whose position was perturbed in relation to the ligand HBD with which it interacts. The 5 spheres within the circle illustrate 5 of the 50 positions occupied by the interacting residue. c) Illustration of how the relative importance of the key ligand atom changes as the distance between it and the perturbed synthetic residue changes. Despite the synthetic residue and ligand atom no longer interacting when the distance between them is greater than 4 Å, the `RF_PLEC_4.5` and `RF_PLEC_5` models continued to rank the ligand atom highly until the synthetic residue-ligand atom distance was greater than the respective PLEC distance cutoff.

### Performance of RF_PLEC models on Contribution dataset

We next examined the performance of the `RF_PLEC` models on our Contribution ZINC dataset. The Contribution generative process more closely approximates real-world protein-ligand binding, as the strength of a synthetic residue-ligand atom interaction is a continuous function of the euclidean distance between them, and typically several high scoring interactions are required in order for a synthetic protein-ligand complex to be deemed active. This is significantly more challenging to learn than the Polar deterministic binding rule, which used a simple distance cutoff and only required a single synthetic residue-ligand atom pair to be ‘active’ for the complex to be active. As with the dataset generated by the Polar generative process, we first fit an `RF_Morgan` model on the Contribution ZINC dataset to assess whether the models were susceptible to ligand-specific bias. The RF Morgan model attained an accuracy of 0.47, suggesting that the structure-based models would have to use the protein to achieve strong predictive performance. All of the `RF_PLEC` models attained a low Spearman’s Rank Correlation Coefficient between the model assigned atom ranks and the ground truth atom ranks, ranging from 0.006 (`RF_PLEC 2.5`) to 0.093 (`(RF_PLEC_6)`). These values suggest that the `RF_PLEC` models were unable to learn the deterministic binding rule. We next assessed whether the models were able to correctly assign a high rank to the most important functional groups. To do this, we computed the Relative Efficiency of Ranking (RER) attained by each `RF_PLEC` model when compared to the ”perfect” rankings obtained by ranking the atoms in a molecule by their ground truth contribution (see Methods). The RER attained by the random baseline was 7.38, and the optimal attainable value for a model is 1.00. The RER scores attained by the `RF_PLEC` models ranged between 4.66 (`RF_PLEC_5`) and 6.68 (`RF_PLEC_2.5`), suggesting that the models struggled to identify the most important functional groups when making predictions. The close-to-random performance of some of the `RF_PLEC` models is also illustrated by Figure 4, which shows the Mean Above Threshold Ranking (MATR) score (see Methods) attained by each model for a variety of thresholds.

**Figure 4:**
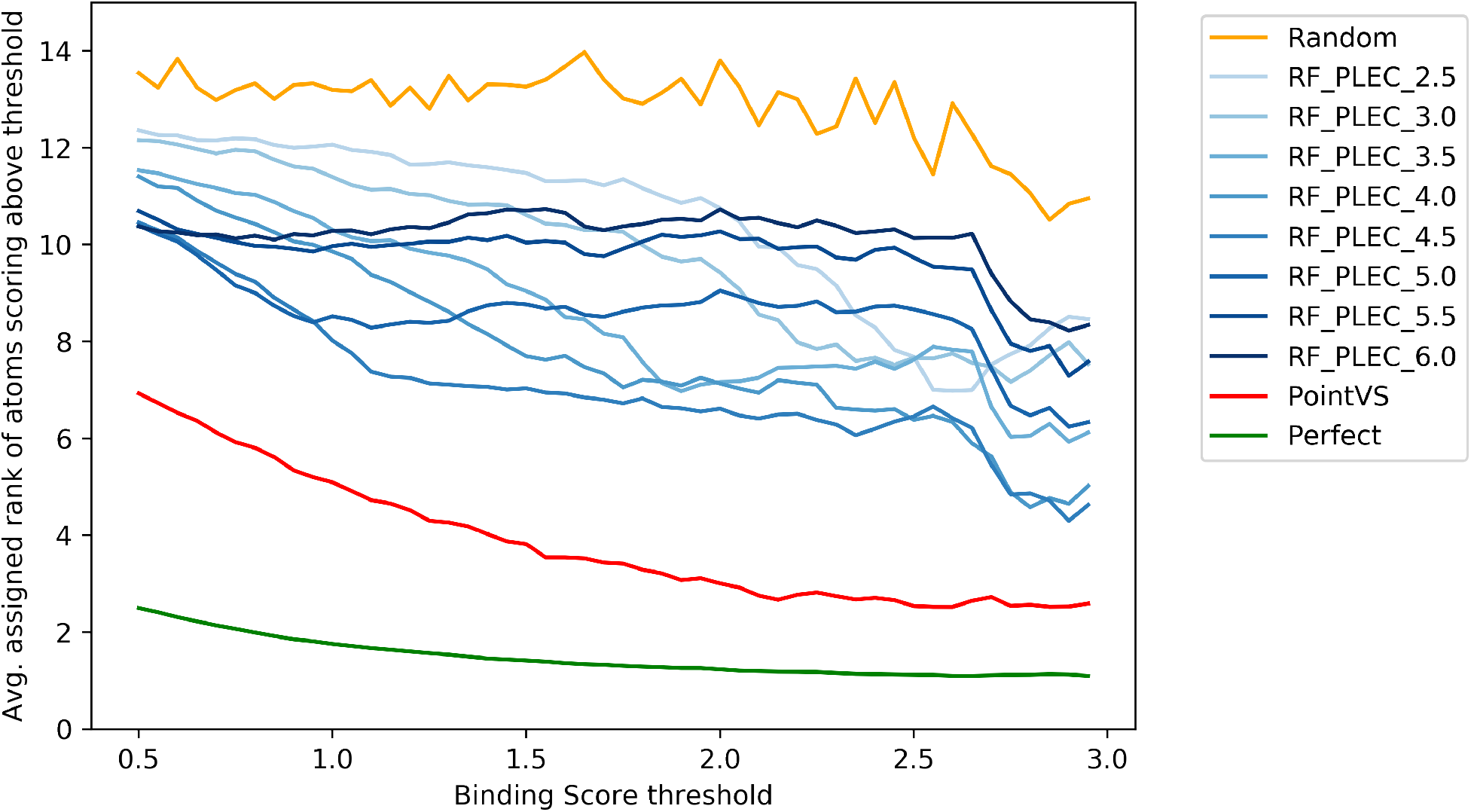
The average rank assigned to all ligand atoms attaining a score above a specified threshold. Perfect is the curve attained when ranking atoms by the true atomic contribution, whilst random is the curve obtained when the atoms are ranked completely at random.

Our results suggest that whilst the PLEC fingerprints were often able to encode a sufficiently detailed level of spatial information for the simplistic Polar tasks, its representation of spatial information is inadequate to learn the more complicated Contribution binding rule. As the Contribution binding rule is itself considerably simpler than the rules which govern real-world protein-ligand binding, it is likely that PLEC (and other fingerprint-based methods which are based on a binary distance cutoff) are ill-equipped to ascertain which functional groups make a key contribution towards binding.

### Quantification of ligand-specific bias on attribution performance

Although the above results suggest that fingerprint-based models struggle to learn complex binding rules, several previous studies (e.g. ^13,33^) have reported that fingerprint-based methods have attained strong predictive accuracy on real-world virtual screening datasets such as DUD-E.^29^ However, recent studies (e.g.^14,30^) have illustrated that virtual screening models are able to attain inflated performance by learning ligand-specific biases, with Sieg et al^30^ reporting that highly simplistic models (e.g. a model with the number of Hydrogen Bond Acceptors in a ligand as the only feature) served as highly accurate classifiers for certain DUD-E targets. Therefore, it is of interest to assess the extent to which realworld factors such as ligand-specific biases and labelling errors degrade the ability of virtual screening models to identify important ligand functional groups.

Although we found that the models trained on DUD-E and LIT-PCBA datasets attained strong predictive accuracy on their respective test sets (Table S2), overall we found that training the models on a dataset which exhibited ligand-specific bias degraded their attribution performance. Table 2 shows the Attribution AUC values attained by the best performing `RF_PLEC` models when trained on a variety of different datasets and tested on the external PDBBind set, generated using the Polar generative process. When training on the DUDE datasets, each of the `RF_PLEC_4` models attained an attribution AUC value marginally below that obtained by the corresponding model trained on the ZINC dataset. However, we observed a considerably larger difference between the attribution AUC values obtained by the `RF_PLEC_4.5` and `RF_PLEC_5` models trained using the DUD-E datasets and the corresponding models trained using the ZINC ligands. Indeed, two of the `RF_PLEC_5` models attained an Attribution AUC (FA10: 0.555, VGFR2: 0.559) which was only marginally higher than the random baseline (0.503). We observed a similar trend for the `RF_PLEC` models trained using LIT-PCBA ligands.

**Table 2:** Attribution AUC obtained by the different `RF_PLEC` models and PointVS on the Polar_ZINC dataset. The best performing dataset for each model is highlighted in **bold**. The `RF_PLEC` models trained on the unbiased ZINC_Polar dataset consistently attained a larger Attribution AUC than those trained on datasets susceptible to ligand-specific bias, suggesting that if a model learns to classify examples based on ligand-specific features, its ability to learn the true binding rule is impaired. By contrast, the highest Attribution AUC attained by PointVS was when training on the LIT-ALDH1 dataset, potentially illustrating that in some instances it was able to learn the deterministic binding rule in the presence of ligand-specific bias.

| Dataset | RF_PLEC_4 | RF_PLEC_4.5 | RF_PLEC_5 | PointVS |
| --- | --- | --- | --- | --- |
| ZINC | **0.89** | **0.86** | **0.78** | 0.85 |
| DUDE-AA2AR | 0.86 | 0.76 | 0.73 | 0.84 |
| DUDE-DRD3 | 0.83 | 0.68 | 0.63 | 0.69 |
| DUDE-FA10 | 0.86 | 0.67 | 0.56 | 0.82 |
| DUDE-MK14 | 0.82 | 0.63 | 0.61 | 0.73 |
| DUDE-VGFR2 | 0.77 | 0.62 | 0.56 | 0.61 |
| LIT-ALDH1 | 0.86 | 0.811 | 0.731 | **0.93** |
| LIT-FEN1 | 0.83 | 0.69 | 0.62 | 0.87 |
| LIT-MAPK1 | 0.81 | 0.61 | 0.59 | 0.70 |
| LIT-PKM2 | 0.81 | 0.73 | 0.67 | 0.82 |
| LIT-VDR | 0.85 | 0.71 | 0.65 | 0.88 |
| Random | 0.503 | 0.503 | 0.503 | 0.503 |

In contrast to the attribution performance attained on the Polar datasets, where the `RF_PLEC_4` models were relatively robust to the introduction of ligand specific bias, the attribution performance attained on the Contribution datasets was substantially degraded in the presence of ligand-specific bias. Figure S1 compares the attribution performance of the `RF_PLEC_4` model when trained on the Contribution ZINC dataset to the models trained on the DUD-E and LIT-PCBA datasets. Training on DUD-E or LIT-PCBA ligands degraded the attribution performance, with the average rank assigned to important ligand atoms decreasing substantially.

### Performance of PointVS

We next examined the performance of an EGNN-based ^27^ method, PointVS (see Methods). When trained using the Polar_ZINC dataset, the PointVS attained an accuracy of 0.886, a AU-PRC of 0.948and an Attribution AUC of 0.851 (Table 1). Whilst PointVS performed slightly worse than the `RF_PLEC_4` and `RF_PLEC_4.5` models, those models were able to exploit the fact that the PLEC featurisation explicitly encoded information about which synthetic residue-ligand atom pairs were near- or below the deterministic binding threshold. By contrast, PointVS was only provided with the unprocessed atomic coordinates, making it more challenging to learn the deterministic binding rule.

When trained and tested on datasets using the Contribution generative process we found that PointVS comfortably outperformed all of the `RF_PLEC` models. Whilst the Spearman’s rank correlation (0.18) obtained by PointVS was not particularly strong, it was higher than any correlation attained by the `RF_PLEC` models (max: `RF_PLEC_6`, 0.09). As with the `RF_PLEC` models, we computed the RER score (see Methods, Evaluation Metrics) to compare PointVS with the ground truth attributions. PointVS attained an RER score of 2.86, considerably better than the top-performing `RF_PLEC` model (`RF_PLEC_5`, 4.66).Figure 4 illustrates the Mean Above Threshold Ranking (MATR) score attained by each model for a variety of thresholds, and shows that PointVS uniformly outperformed all `RF_PLEC` models for all thresholds. These results are perhaps not surprising, as the `RF_PLEC` models do not capture a precise encoding of each atom’s position, only recording whether the distance between two atoms is below a pre-specified threshold, making it difficult for the `RF_PLEC` models to approximate the nonlinear deterministic binding rule. By contrast, PointVS is not constrained by a particular featurization and can learn the deterministic binding rule from the data.

When trained on the DUD-E or LIT-PCBA datasets, PointVS demonstrated a degree of robustness to ligand-specific bias under the Polar generative process (Table 2), although in several cases the Attribution was substantially lower than the value obtained for the Polar_ZINC data. Under the Contribution generative process, consistent with the results obtained using the `RF_PLEC_4` model, PointVS’ attribution performance was considerably worse when trained with the DUD-E or LIT-PCBA datasets compared to the Contribution ZINC dataset (Figure 5). As both the PointVS and `RF_PLEC_4` models struggle to learn complex binding rules in the presence of ligand-specific bias, it is likely that the models would be unable to identify the functional groups responsible for binding on real-world datasets if similar biases were present.

**Figure 5:**
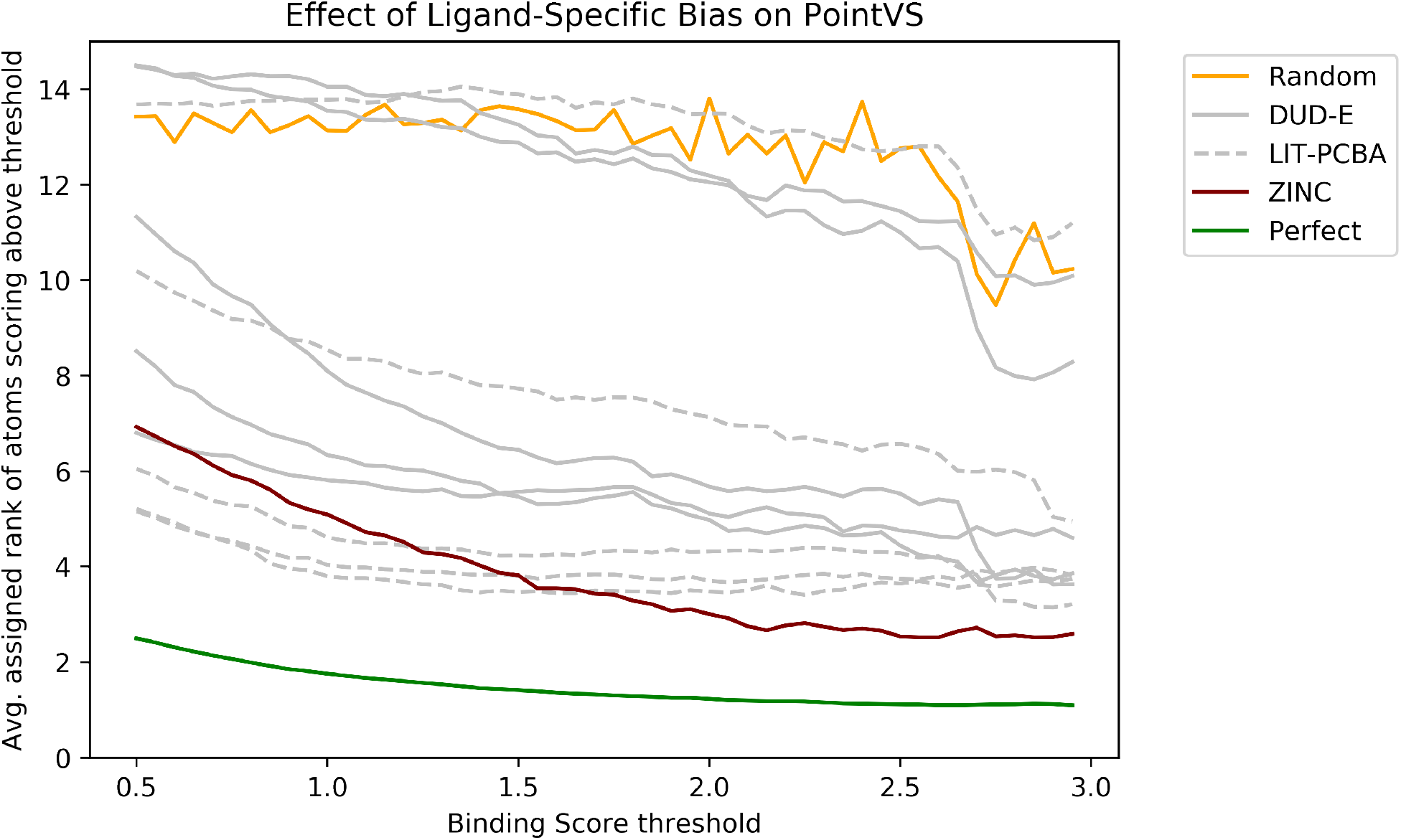
The average rank assigned to all ligand atoms attaining a score above a specified threshold on the PDBBind test set. Each line corresponds to the performance obtained by a PointVS model trained on a different training set. The model trained on the unbiased ZINC dataset outperforms the models trained on the real-world DUD-E and LIT-PCBA datasets, illustrating that ligand-specific biases hamper the ability of virtual screening models to identify the most important functional groups.

## Conclusion

We have proposed a framework for assessing the ability of different ML algorithms to attribute binding at the atomic level. By simulating synthetic datasets we are able to define a ground truth for which atoms in a synthetic protein-ligand complex are responsible for binding, something which is not currently possible with real-world data. Despite several recent studies suggesting that fingerprint-based machine learning algorithms achieve comparable performance with current deep learning methods, we found that a deep learning-based EGNN was better able to elucidate which atoms were responsible for ‘binding’ when the training datasets were not susceptible to ligand-specific biases. This suggests that deep learning methods should have a greater ability to generalize to novel targets.

When we trained our models on datasets containing ligand-specific bias, we found that a model’s ability to accurately assign atomic importance was diminished, consistent with the notion that dataset-specific bias degrades model generalizability.

However, for the Polar datasets, we found that the drop in attribution performance attained by PointVS was far smaller than the corresponding drop in performance observed for the `RF_PLEC_4.5` and `RF_PLEC_5` models and similar to that observed for the `RF_PLEC_4` model, suggesting that models which can more easily learn the true binding rule are less susceptible to the effects of bias. The development of models which are able to capture and apply important spatial information should therefore continue to be a priority.

Our analysis has a number of limitations. The deterministic binding rules used in both the Polar and Contribution generative processes are significantly simplified compared to real-world protein-ligand binding. However, our objective was to assess the extent to which different ML algorithms were able to capture and apply relevant spatial information and our relatively simple generative processes were sufficient to highlight the difficulties of explicitly encoding a precise level of 3D information into a fingerprint.

Whilst there is significant interest in developing models which can predict protein-ligand binding with high accuracy, there is a strong need for models which can identify the functional groups which contribute most heavily towards binding; the identification of key functional groups would allow easier iterative optimisation of lead compounds. Moreover, models which demonstrated an understanding of biophysical rules would be more likely to be accepted by human experts. In addition to the development of machine learning architectures which can better capture and apply important spatial information, avenues of research which might better enable models to identify important function groups include data augmentation and multi-task learning. An alternative strategy which can improve the quality of virtual screening models is the incorporation of experimentally verified misses into the training data.^34^

We hope that our synthetic framework will prove useful to researchers seeking to benchmark the ability of different virtual screening models to capture and apply important spatial information. We have made the datasets used in this study available, alongside the code to generate synthetic protein-ligand complexes from a set of ligands.

## Code and Data Availability

Code is available at https://github.com/tomhadfield95/synthVS. Datasets are available at https://opig.stats.ox.ac.uk/data/downloads/synthVS_data_for_release.tar.gz

## Declarations

T.E.H. is an employee of AstraZeneca PLC; all work was done whilst a doctoral student at the University of Oxford. C.M.D. is an employee of Exscientia PLC.

## Acknowledgements

T.E.H. is supported by funding from the Engineering and Physical Sciences Research Council (EPSRC), LifeArc, F. Hoffmann-La Roche AG, and UCB Pharma (Reference: EP/L016044/1).

J.S. is supported by funding from the Biotechnology and Biosciences Research Council (BB-SRC) BB/S507611/1 and BenevolentAI..

The authors would like to thank Fergus Boyles, Kris Birchall and Andy Merritt for helpful discussions.

