## Supporting Information for "Exploring The Ability Of Machine Learning-Based Virtual Screening Models To Identify The Functional Groups Responsible For Binding"

### Supporting Information
### Exploring The Ability Of Predictive Models To Identify The Functional Groups Responsible For Protein-Ligand Binding

Thomas E. Hadfield, Jack Scantlebury, and Charlotte M. Deane*

*Oxford Protein Informatics Group, Department of Statistics, University of Oxford, Oxford OX1 3LB, UK*

Table S1: Performance of the `RF_Morgan` model on different datasets. Predictive accuracy substantially better than random suggests that the datasets may suffer from ligand-specific bias. Accuracy denotes the proportion of correctly classified examples, whereas Random Accuracy denotes the accuracy that would have been obtained by assigning all examples the most common label (= max(% actives, % inactives). AU-PRC denotes the area under the Precision-Recall curve. Balanced Accuracy and Balanced AU-PRC denote the respective accuracy and area under the Precision-Recall curve when the model was trained using an equivalent number of actives and inactives.

| Dataset | Random Accuracy | Accuracy | AU-PRC | Balanced Accuracy | Balanced AU-PRC |
|---|---|---|---|---|---|
| ZINC | 0.504 | 0.52 | 0.53 | N/A | N/A |
| DUDE-AA2AR | 0.93 | 1.0 | 1.0 | 0.984 | 0.996 |
| DUDE-DRD3 | 0.972 | 0.998 | 1.0 | 0.978 | 0.995 |
| DUDE-FA10 | 0.97 | 1.0 | 1.0 | 0.992 | 1.0 |
| DUDE-MK14 | 0.976 | 0.998 | 1.0 | 0.994 | 1.0 |
| DUDE-VGFR2 | 0.984 | 1.0 | 1.0 | 0.99 | 0.999 |
| LIT-ALDH1 | 0.613 | 0.76 | 0.809 | 0.768 | 0.806 |
| LIT-FEN1 | 0.956 | 0.958 | 0.584 | 0.778 | 0.883 |
| LIT-MAPK1 | 0.964 | 0.964 | 0.292 | 0.692 | 0.797 |
| LIT-PKM2 | 0.944 | 0.952 | 0.755 | 79 | 0.901 |
| LIT-VDR | 0.928 | 0.942 | 0.6 | 0.772 | 0.87 |

Table S2: Performance of the `RF_PLEC_4` model on different datasets. Accuracy denotes the proportion of correctly classified examples, whereas Random Accuracy denotes the accuracy that would have been obtained by assigning all examples the most common label (= max(% actives, % inactives). AU-PRC denotes the area under the Precision-Recall curve. Balanced Accuracy and Balanced AU-PRC denote the respective accuracy and area under the Precision-Recall curve when the model was trained using an equivalent number of actives and inactives.

| Dataset | Random Accuracy | Accuracy | AU-PRC | Balanced Accuracy | Balanced AU-PRC |
|---|---|---|---|---|---|
| ZINC | 0.504 | 0.948 | 0.988 | N/A | N/A |
| DUDE-AA2AR | 0.93 | 0.992 | 0.987 | 0.942 | 0.992 |
| DUDE-DRD3 | 0.972 | 0.992 | 0.974 | 0.904 | 0.981 |
| DUDE-FA10 | 0.97 | 0.992 | 0.971 | 0.934 | 0.993 |
| DUDE-MK14 | 0.976 | 0.99 | 0.994 | 0.928 | 0.985 |
| DUDE-VGFR2 | 0.984 | 0.992 | 0.92 | 0.926 | 0.987 |
| LIT-ALDH1 | 0.613 | 0.948 | 0.987 | 0.946 | 0.988 |
| LIT-FEN1 | 0.956 | 0.968 | 0.839 | 0.858 | 0.948 |
| LIT-MAPK1 | 0.964 | 0.972 | 0.672 | 0.822 | 0.923 |
| LIT-PKM2 | 0.944 | 0.974 | 0.976 | 0.916 | 0.984 |
| LIT-VDR | 0.928 | 0.98 | 0.931 | 0.928 | 0.979 |

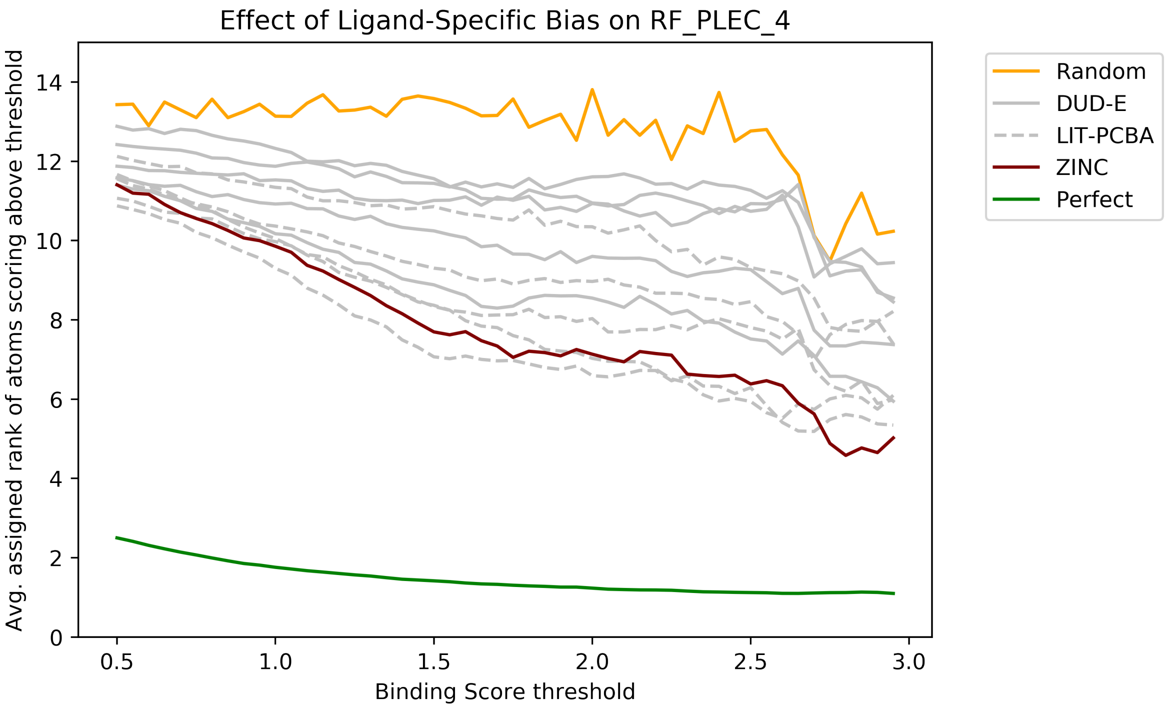


Figure S1: The average rank assigned to all ligand atoms attaining a score above a specified threshold on the PDBBind test set. Each line corresponds to the performance obtained by an `RF_PLEC_4` model trained on a different training set. The model trained on the unbiased ZINC dataset outperforms the models trained on the real-world DUD-E and LIT-PCBA datasets, illustrating that ligand-specific biases hamper the ability of virtual screening models to identify the most important functional groups.
